# The biogeography of the Mediterranean Taurus Mountains: Illustrating how complex topography and climatic conditions have shaped their biodiversity

**DOI:** 10.1101/2021.09.15.460441

**Authors:** Hakan Gür

## Abstract

Climate is well known as the main driver of species distributions. In this study, I focused on Geyik Mountains and surrounding areas to illustrate how complex topography and climatic conditions have shaped the distribution patterns of species/communities and therefore the biodiversity of the Mediterranean Taurus Mountains, one of the most biologically diverse areas in the Mediterranean Basin biodiversity hotspot. Accordingly, I used an ecological niche modelling approach, which has been widely used in recent biogeographic studies. I chose Taurus ground squirrels (*Spermophilus taurensis*) and coniferous forests as the representatives of high- and low-altitude species/communities, respectively. The results simply illustrate how complex topography and temperature and precipitation gradients have had a substantial role in shaping the distribution patterns of species/communities and therefore the biodiversity of the Mediterranean Taurus Mountains.

## Background

A region should meet two main criteria to be identified as a biodiversity hotspot. A biodiversity hotspot should have (1) ≥ 1500 endemic vascular plant species and (2) ≤ 30% of its original natural vegetation. In other words, it should be irreplaceable and threatened. According to these criteria, 36 biodiversity hotspots were identified around the world. The forests and other remnant habitats in these biodiversity hotspots correspond to only 2.4% of Earth’s land surface, but still have more than 50% of endemic plant species and nearly 43% of endemic amphibian, reptile, bird, and mammal species in the world (Conservation International, 2019).

Anatolia (the Asian part of Turkey) is geologically located in the Alpine-Himalayan orogenic belt (Mather, 2009) and the region where three of the world’s 36 biodiversity hotspots meet, and interact: the (1) Caucasus, (2) Irano-Anatolian, and (3) Mediterranean Basin biodiversity hotspots. This means that Anatolia has a high biodiversity and a high percentage of life found nowhere else in the world, but has lost most of its original natural vegetation. In other words, Anatolia is one of the world’s biologically richest and most threatened terrestrial regions (Conservation International, 2019; see also Gür, 2016, 2017a, 2017b).

The Mediterranean Basin biodiversity hotspot is one of the most important biodiversity hotspots in the world, with high levels of endemism, particularly for plant species (Conservation International, 2019; see also Özüdoğru et al., 2020), and in Anatolia, includes four ecoregions (“large unit(s) of land or water containing a geographically distinct assemblage of species, natural communities, and environmental conditions”; World Wildlife Fund, 2019): (1) ‘Southeastern Europe: along the coastline of Greece and Turkey, stretching into Macedonia’ (PA1201), (2) ‘Southeastern Europe: Western Turkey’ (PA1202), (3) ‘Southwestern Asia: along the coast of the Mediterranean Sea in Turkey, Jordan, Israel, and Syria’ (PA1207), and (4) ‘Western Asia: Southern Turkey into Syria, Lebanon, Israel’ (PA1220) (Figure 1). All these ecoregions have Critical/Endangered status, and are from the ‘Mediterranean forests, woodlands, and scrubs’ habitat type, one of the world’s 14 major terrestrial habitat types that “share similar environmental conditions, habitat structure, and patterns of biological complexity, and that contain similar communities and species adaptations” (World Wildlife Fund, 2019). This habitat type is characterized by hot and dry summers and cool and wet winters (World Wildlife Fund, 2019).

**Figure 1.**
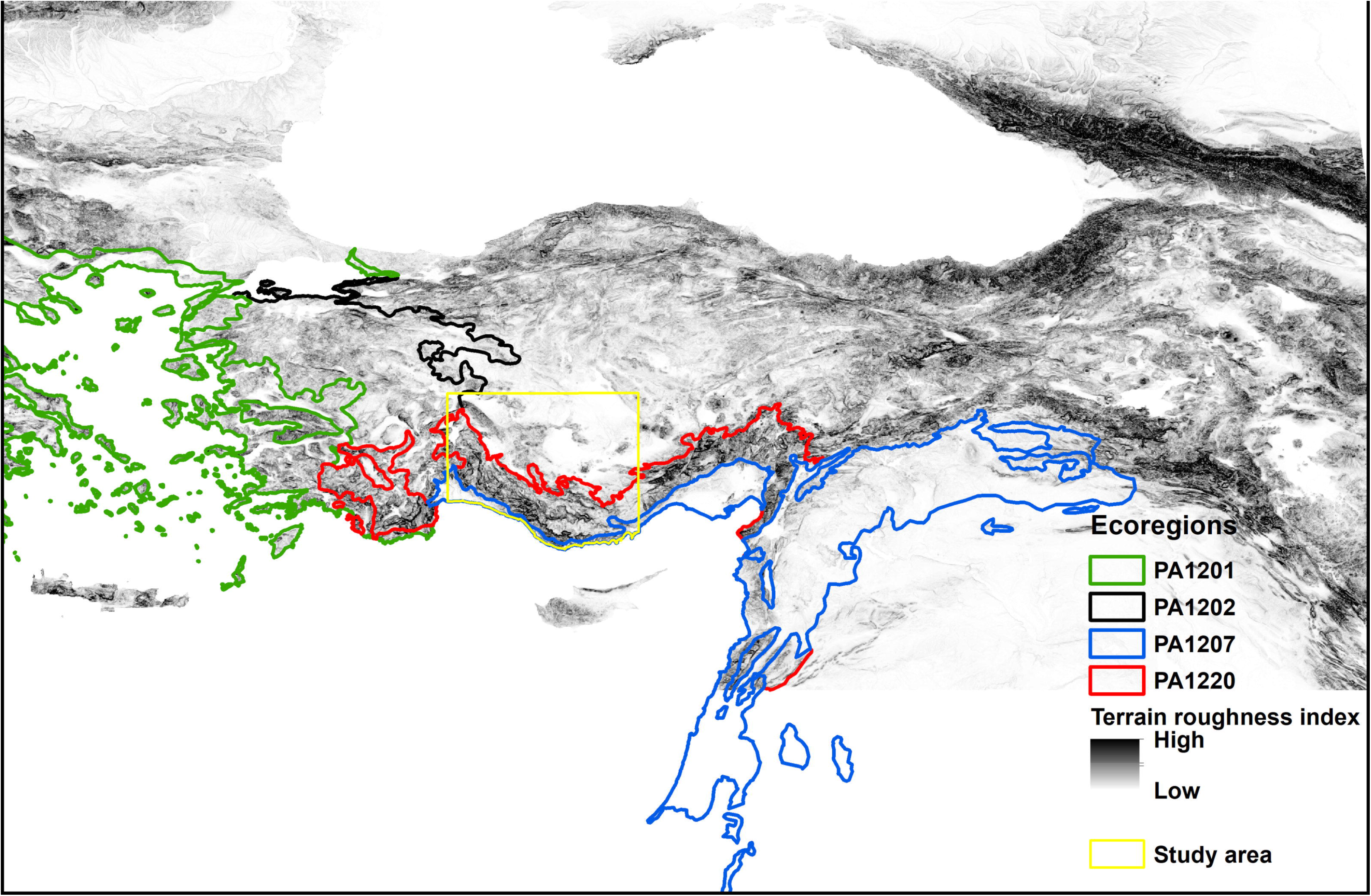
The ecoregions of the Mediterranean Basin biodiversity hotspot and terrain roughness in Anatolia. PA1201: ‘Southeastern Europe: along the coastline of Greece and Turkey, stretching into Macedonia’, PA1202: ‘Southeastern Europe: Western Turkey’, PA1207: ‘Southwestern Asia: along the coast of the Mediterranean Sea in Turkey, Jordan, Israel, and Syria’, and PA1220: ‘Western Asia: Southern Turkey into Syria, Lebanon, Israel’. The ecoregions were drawn based on World Wildlife Fund (2019), and terrain roughness index is from ENVIREM database (Title & Bemmels, 2017).

The ‘Western Asia’ ecoregion (PA1220) is one of the most biologically diverse areas in the Mediterranean Basin biodiversity hotspot (Blondel & Médail, 2009; Kryštufek, Vohralík, & Obuch, 2009; Médail & Diadema, 2009; Médail & Quézel, 1997; Parolly, 2015; World Wildlife Fund, 2019), and in Anatolia, includes the western and central Taurus Mountains located along the Mediterranean coast of Anatolia (hereafter referred to as the Mediterranean Taurus Mountains; Figure 2). These mountain ranges have complex topography (Figure 1 and 2) and climatic conditions (Atalay, Efe, & Öztürk, 2014; Gür, 2017b; Parolly, 2015), and have acted as refugia during Quaternary glacial-interglacial cycles (Médail & Diadema, 2009; see also Gür, Perktaş, & Kart Gür, 2018) because only their peaks were glaciated (Sarıkaya & Çiner, 2017). These main factors have contributed to high levels of species diversity and endemism (Allen, 2009; Blondel, Aronson, Bodiou, & Boeuf, 2010; Médail & Diadema, 2009; Thompson, 2005; World Wildlife Fund, 2019). The overlapping of the Mediterranean and Irano-Turanian floristic zones (or the Mediterranean Basin and Irano-Anatolian biodiversity hotspots or the ecoregions from the ‘Mediterranean forests, woodlands, and scrubs’ and ‘Temperate grasslands, savannas and shrublands’ major habitat types) in these mountain ranges has also contributed to the evolution of unique biodiversity (Parolly, 2004; World Wildlife Fund, 2019; Figure 2). Despite their importance with respect to biodiversity, the Mediterranean Taurus Mountains are an under-explored region of the Mediterranean Basin biodiversity hotspot (Médail & Diadema, 2009). Thus, I believe that future studies, based on the key approaches widely used in recent biogeographic studies (e.g. see Gür, 2019), are really needed to accurately understand the biodiversity of these mountain ranges.

**Figure 2.**
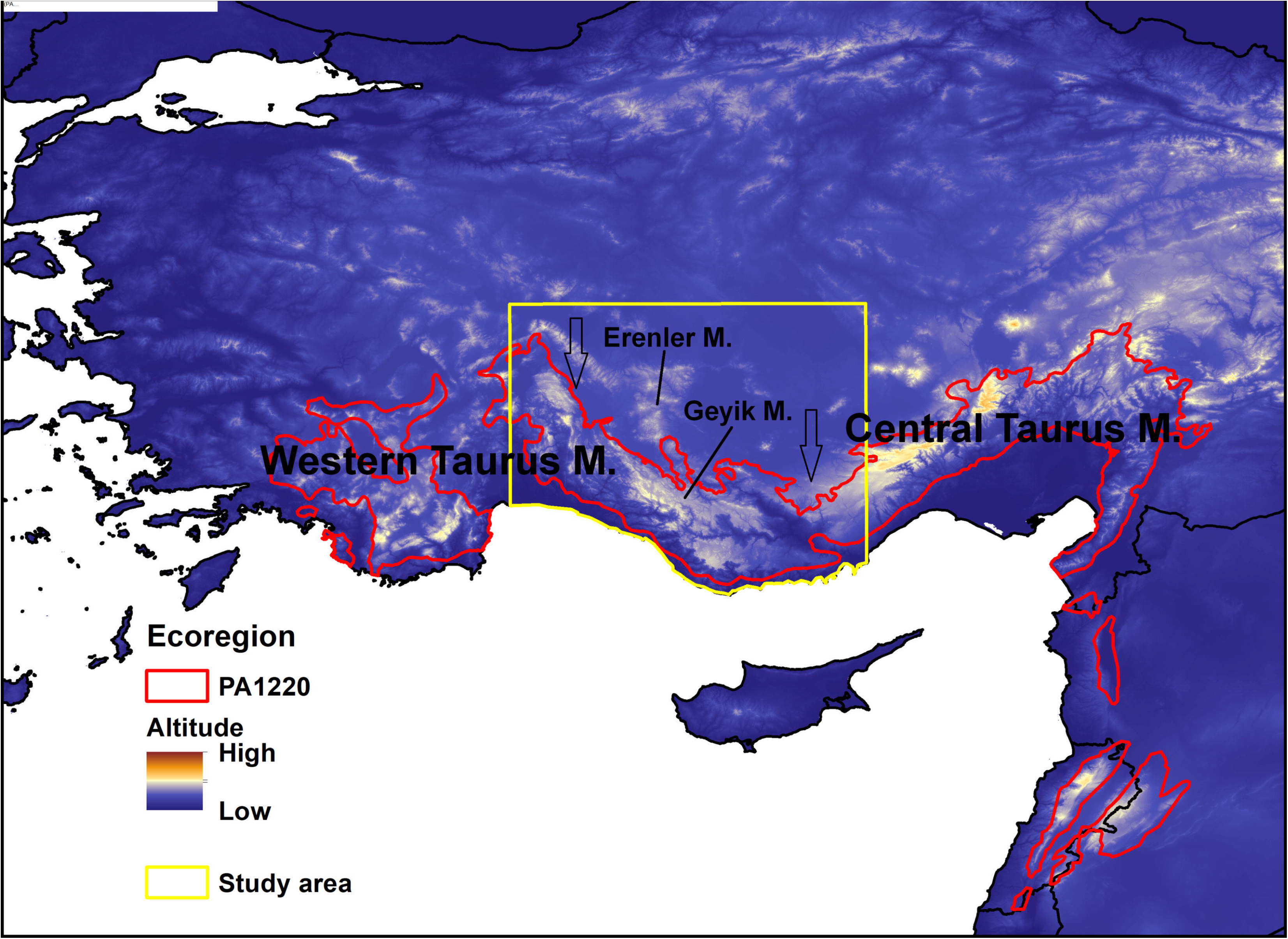
The ‘Western Asia: Southern Turkey into Syria, Lebanon, Israel’ ecoregion (PA1220) and altitude in the southern Anatolia. The arrows represent the influence of the Irano-Turanian floristic zone. The ecoregion was drawn based on World Wildlife Fund (2019), and altitude is from Jarvis, Reuter, Nelson, & Guevara (2008).

Climate is well known as the main driver of species distributions. Thus, a key approach to understanding the distribution patterns of species and therefore the biogeography of a region is to use ecological niche modelling. Ecological niche modelling relates georeferenced species occurrence data (a set of geographical coordinates where species of interest has been observed) to environmental data (a number of environmental variables obtained using a geographical information system-based approach), and creates models to predict the distribution patterns of species (Peterson & Anamza, 2015; Peterson et al., 2011).

### A case study

In this study, I focused on Geyik Mountains and surrounding areas (Figure 1 and 2) to illustrate how complex topography and climatic conditions have shaped the distribution patterns of species/communities and therefore the biodiversity of the Mediterranean Taurus Mountains, one of the most biologically diverse areas in the Mediterranean Basin biodiversity hotspot. Accordingly, I used an ecological niche modelling approach, which has been widely used in recent biogeographic studies (Perktaş & Gür, 2015; Peterson & Anamza, 2015; Peterson et al., 2011). Thus, I also exemplified how this approach can be used to understand the biogeography of a region. It is important to note here that historical factors have doubtless had a substantial role in shaping the biodiversity of these mountain ranges (e.g. Thompson, 2005). Gür et al. (2018) have already studied one of these historical factors, the refugial role of these mountain ranges by examining how global climate changes through the Late Quaternary glacial-interglacial cycles have affected high-altitude species/communities (in particular, Taurus ground squirrels, *Spermophilus taurensis*). Taurus ground squirrels are group-living, diurnal, hibernating, and pre-dominantly herbivorous, burrowing ground squirrels. They are endemic to a small range, i.e. to Erenler Mountain in the north and Geyik Mountains in the south. They inhabit mainly grasslands at the higher altitudes, e.g. > ~ 1500 m in the sea-facing side of Geyik Mountains (Gür et al., 2018 and references therein), being located in the subalpine herbaceous vegetation belt (Atalay et al., 2014). The sea-facing side of the Mediterranean Taurus Mountains also harbors mainly coniferous forests at the lower altitudes. These forests are located in two vegetation belts: (1) the Eu-Mediterranean belt, with Calabrian pine (*Pinus brutia*) as the dominant coniferous species, and (2) the Oro-Mediterranean belt, with Anatolian black pine (*Pinus nigra* subsp. *pallasiana*), Lebanon cedar (*Cedrus libani*), Taurus fir (*Abies cilicica*), and junipers (*Juniperus foettidissima* and *J. excelsa*) as the dominant coniferous species (Atalay et al., 2014; Kaya & Raynal, 2001; World Wildlife Fund, 2019). Thus, I chose Taurus ground squirrels and coniferous forests as the representatives of high- and low-altitude species/communities, respectively. Ecological niche modelling has been previously used successfully to model the distribution patterns of both Taurus ground squirrels (Gür et al., 2018) and forests of the other regions (e.g. Carnaval & Moritz, 2008; Werneck, Costa, Colli, Prado, & Sites, 2011)

### A short methodology

Occurrence records for Taurus ground squirrels (*Spermophilus taurensis*) were obtained from Gür et al. (2018) and additional field studies (Figure 3). These records were spatially filtered by reducing multiple records to a single record within 4 km distance, resulting in 50 records used for ecological niche modelling.

**Figure 3.**
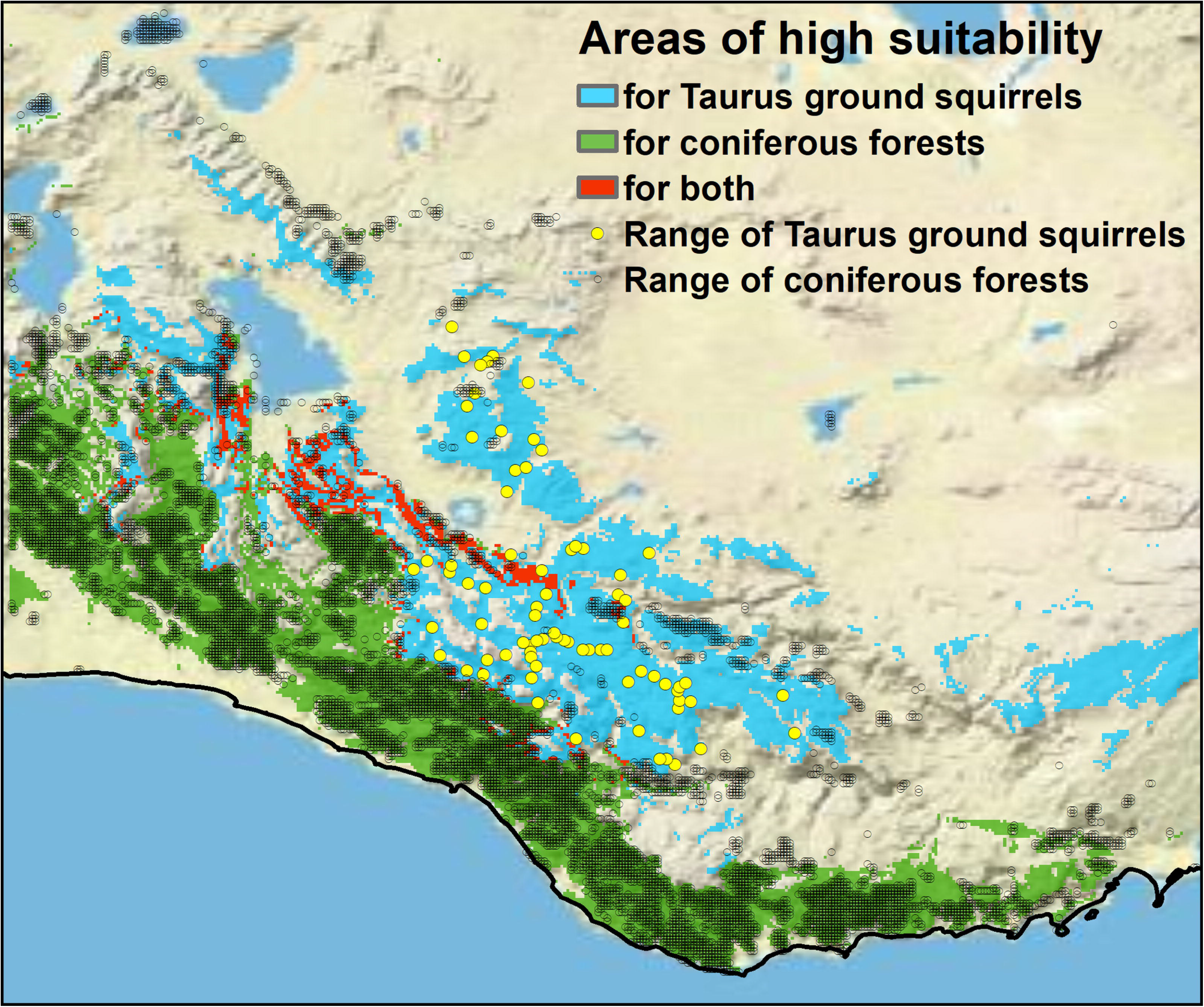
Areas of high suitability for Taurus ground squirrels (*Spermophilus taurensis*), for coniferous forests, and for both of these. The ranges of Taurus ground squirrels and coniferous forests are represented by the occurrence records (N = 76 and 8644, respectively).

Occurrence records for coniferous forests were obtained from a raster version, subsequently resampled to a spatial resolution of 30 arc-seconds, of the GlobCover 2009 land cover map V2.3, processed by the European Space Agency (http://due.esrin.esa.int/page_globcover.php) (Figure 3). Of raster cells in which the ‘closed (>40%) needle-leaved evergreen forest (>5m)’ land cover type is dominant, 25% (2161 records) were randomly selected to serve as records used for ecological niche modelling.

Bioclimatic data were downloaded from the CHELSA database at a spatial resolution of 30 arc-seconds for present (1979-2013) conditions (Karger et al., 2017a, 2017b). A subset of eight bioclimatic variables was chosen: annual mean temperature and precipitation (BIO1 and 12), temperature and precipitation seasonality (BIO4 and 15), and mean temperature and precipitation of the warmest and coldest quarters (BIO10, 11, 18, and 19) (see also Table 1). All these variables were masked to include Geyik Mountains and surrounding areas (i.e. the study area; 30.76° to 34.07° E and 36.03° to 38.75° N; Figure 1 and 2).

**Table 1.** Statistics of altitude and bioclimatic variables, based on all the grid cells (N) falling in the study area and areas of high suitability for Taurus ground squirrels (*Spermophilus taurensis*) and coniferous forests.

| Variables | Study area (N = 110885) |  |  |  |  | Taurus ground squirrels (N = 11681) |  |  |  |  | Coniferous forests (N = 14580) |  |  |  |  |
| --- | --- | --- | --- | --- | --- | --- | --- | --- | --- | --- | --- | --- | --- | --- | --- |
|  | Mean | SD | Median | Min | Max | Mean | SD | Median | Min | Max | Mean | SD | Median | Min | Max |
| Altitude | 1151 | 440 | 1091 | -3 | 2964 | 1690 | 200 | 1686 | 1229 | 2375 | 813 | 456 | 776 | 15 | 1901 |
| Bio1 | 12.3 | 2.8 | 12.4 | 1.5 | 20.6 | 9.2 | 1.1 | 9.2 | 5.7 | 11.6 | 14.8 | 3.0 | 15.1 | 8.6 | 19.8 |
| Bio4 | 8.0 | 0.4 | 8.1 | 6.5 | 8.5 | 7.9 | 0.2 | 7.9 | 6.9 | 8.3 | 7.6 | 0.4 | 7.8 | 6.5 | 8.1 |
| Bio10 | 23.8 | 2.7 | 24.0 | 13.0 | 31.0 | 20.7 | 1.2 | 20.7 | 17.0 | 23.4 | 25.7 | 2.7 | 25.9 | 19.0 | 30.5 |
| Bio11 | 1.9 | 2.9 | 1.7 | -8.1 | 11.7 | -0.9 | 1 | -0.9 | -4.0 | 1.2 | 5.1 | 3.2 | 5.4 | -1.6 | 11.1 |
| Bio12 | 562 | 242 | 498 | 255 | 1511 | 723 | 151 | 702 | 418 | 1361 | 899 | 169 | 887 | 505 | 1413 |
| Bio15 | 60 | 17 | 53 | 30 | 99 | 70 | 9 | 71 | 46 | 96 | 83 | 8 | 84 | 34 | 99 |
| Bio18 | 22 | 11 | 19 | 3 | 69 | 24 | 8 | 25 | 6 | 42 | 18 | 10 | 19 | 3 | 63 |
| Bio19 | 252 | 150 | 209 | 80 | 821 | 360 | 83 | 349 | 163 | 682 | 468 | 88 | 464 | 129 | 807 |

BIO1: annual mean temperature (°C), BIO4: temperature seasonality (standard deviation, SD), BIO10: mean temperature of the warmest quarters (°C), BIO11: mean temperature of the coldest quarters (°C), BIO12: annual precipitation (mm), BIO15: precipitation seasonality (coefficient of variation), BIO18: precipitation of the warmest quarters (mm), BIO19: precipitation of the coldest quarters (mm).

To model the distribution patterns of Taurus ground squirrels and coniferous forests, the maximum entropy machine learning algorithm in the software MaxEnt version 3.3.3k (Elith et al., 2011; Phillips, Anderson, & Schapire, 2006) was used. MaxEnt was run with the settings described in Gür et al. (2018), except for the cases noted below. The whole study area was used to randomly sample 10000 background points. Single-variable response curves were created for the bioclimatic variables. The contribution of each variable to the best model was determined using a jackknife test (with only one variable). A total of 125 different models were tested based on different model parameters [five feature class types (L, LQ, H, LQH, and LQHPT) and a range of regularization multipliers (1 to 5 in increments of 1)] and training/test partitions (spatial jackknifing, k = 5). All these analyses were conducted using the software SDMtoolbox version 1.1c (Brown, 2014) and the software ArcGIS version 10.2.2.

For Taurus ground squirrels and coniferous forests, an estimate of niche breadth was calculated by applying a traditional measure of niche breadth (inverse concentration; Levin, 1968) to the prediction of bioclimatic suitability for each species/community (i.e. Taurus ground squirrels and coniferous forests, respectively). This analysis was conducted using the software ENMTools version 1.4.4 (Warren, Glor, & Turelli, 2010).

## Results and Discussion

For Taurus ground squirrels, the best model used feature classes of linear, quadratic, and hinge (i.e. LQH) and a regularization multiplier of 1, and performed better than a random prediction (training AUC = 0.949). Areas of high suitability (an equal training sensitivity and specificity logistic threshold value of 0.324 or higher) were predicted mainly at the higher altitudes of Erenler Mountain and Geyik Mountains (Figure 3 and 4), i.e. in areas influenced by both the Mediterranean climate (hot and dry summers and cool and wet winters) and the continental climate (hot and dry summers and cold and snowy winters) (Table 1; for the response curves, see also Figure 5). Areas of high suitability around Beyşehir Lake and in the east represent areas currently unoccupied due to non-climate-related factors or populations not yet detected (Figure 3). Within areas of high suitability, the most common land cover type (47%) is by far the ‘mosaic vegetation (grassland/shrubland/forest) (50-70%) / cropland (20-50%)’. Temperature variables dominated over precipitation variables, suggesting that temperature (or factors associated with temperature) is a more important driver of the distribution patterns of Taurus ground squirrels. Temperature was also identified as the main determinant for the occurrence of high-montane species of different taxonomic groups (fern, vascular plant, wood-inhabiting fungus, mollusk, saproxylic beetle, and bird; Bässler et al., 2010). Gür et al. (2018) suggested that high winter temperature and/or low winter precipitation limit the geographic distribution of Taurus ground squirrels by changing snow cover and/or the vegetation structure. Indeed, the results also support this idea (see below).

**Figure 4.**
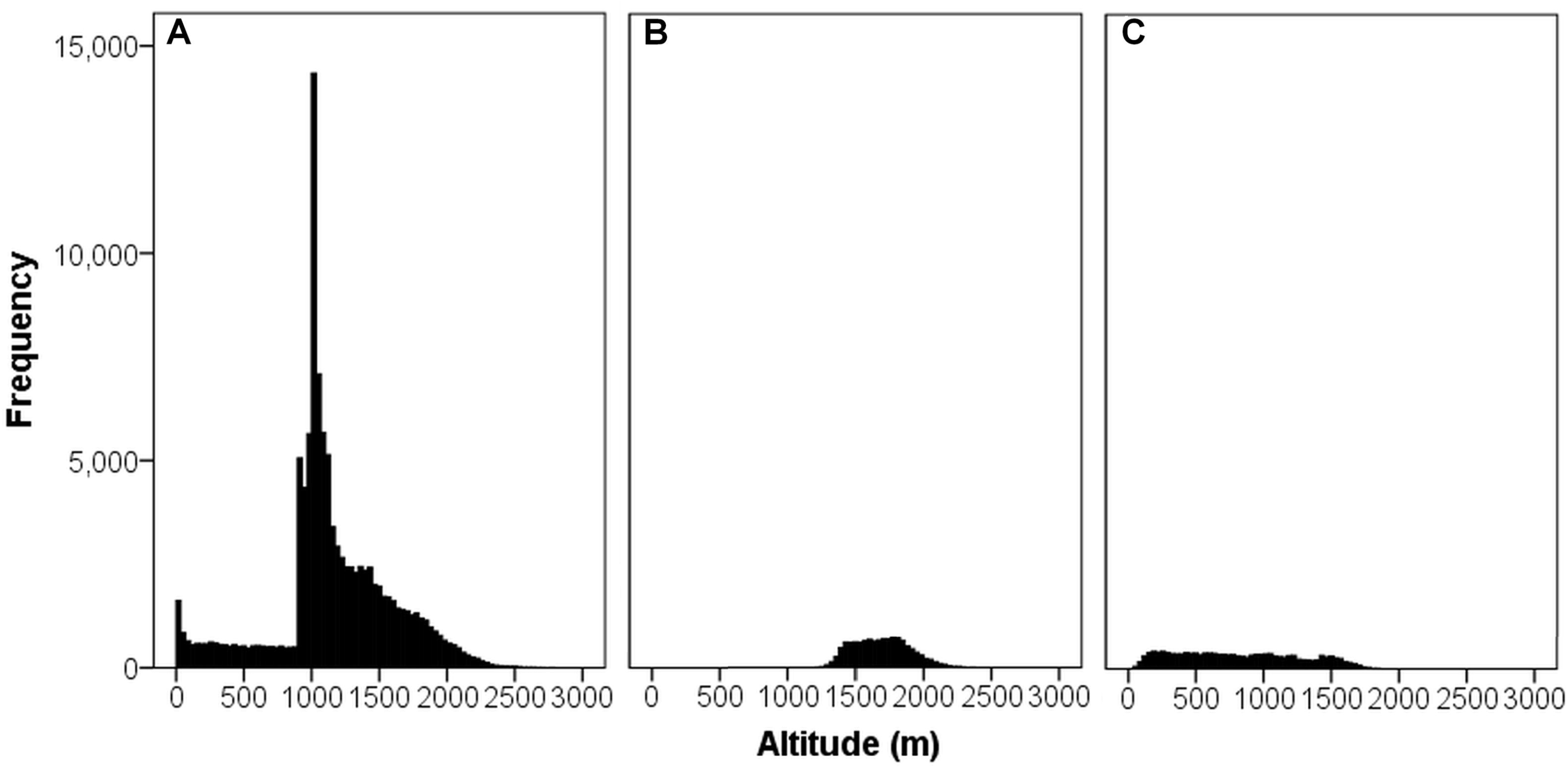
The frequency distribution of altitude, based on all the grid cells falling in (A) the study area (N = 110885) and areas of high suitability for (B) Taurus ground squirrels (*Spermophilus taurensis*) (N = 11681) and (C) coniferous forests (N = 14580).

**Figure 5.**
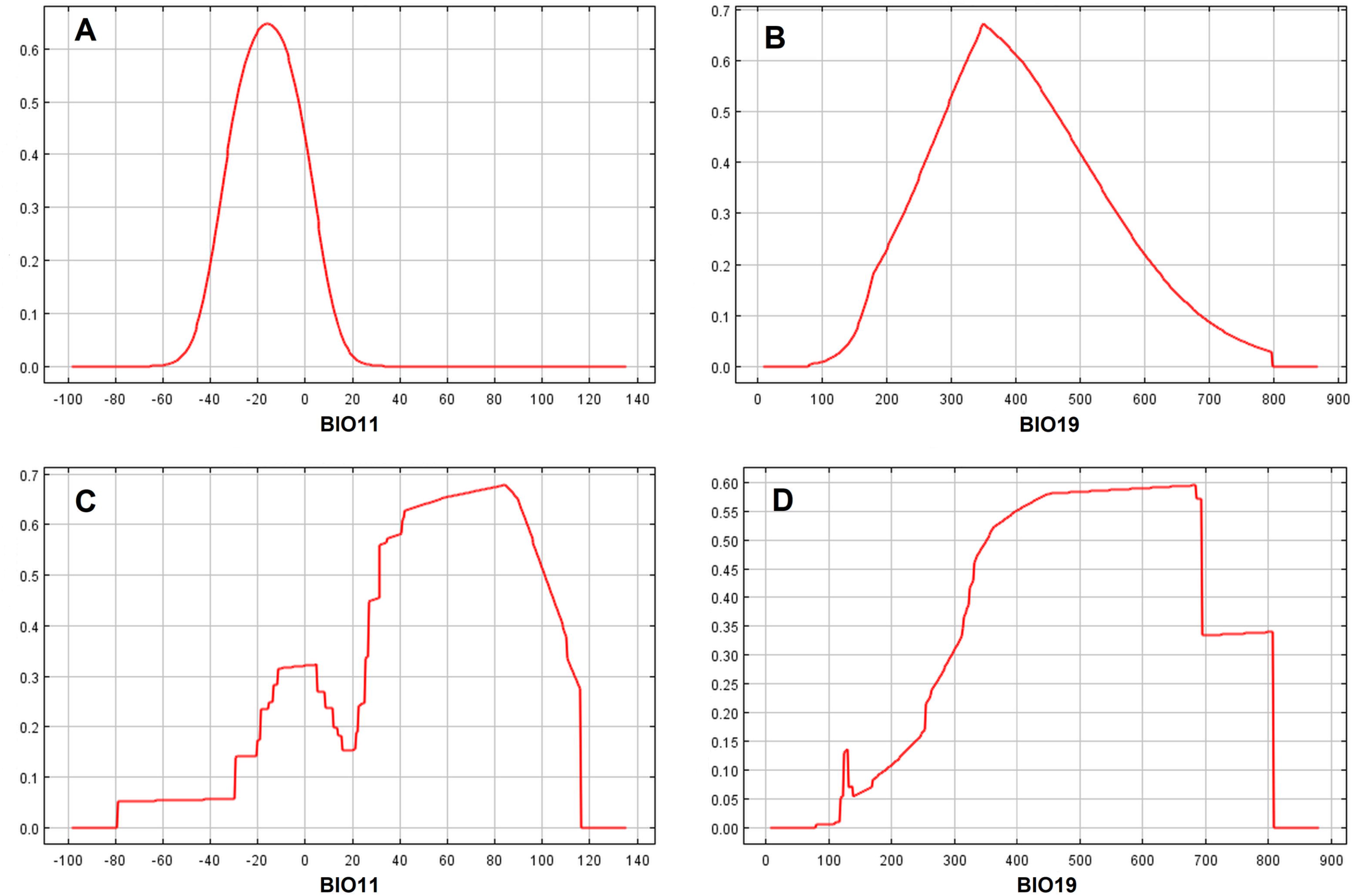
The single variable response curves of (A, C) winter temperature (BIO11) and (B, D) precipitation (BIO19) for (A, B) Taurus ground squirrels (*Spermophilus taurensis*) and (C, D) coniferous forests. Winter temperature is the bioclimatic variable with the highest gain when used in isolation for Taurus ground squirrels, and winter precipitation for coniferous forests.

For coniferous forests, the best model used feature classes of linear, quadratic, hinge, product, and threshold (i.e. LQHPT) and a regularization multiplier of 1, and performed better than a random prediction (training AUC = 0.842). Areas of high suitability (0.442 or higher) were predicted mainly at the lower altitudes in the sea-facing side of the Mediterranean Taurus Mountains (Figure 3 and 4), i.e. in areas having more typical characteristics of the Mediterranean climate (Table 1; for the response curves, see also Figure 5). However, coniferous forests do not occur everywhere within areas of high suitability (Figure 3). This may be due to smaller-scale variations in the distribution patterns (see below) and/or modelling errors (e.g. those associated with the occurrence records obtained from the land cover map). However, this may also be a result of the reality that “the Mediterranean Basin has experienced intensive human development and impact on its ecosystems for thousands of years, significantly longer than any other hotspot” (Conservation International, 2019). Indeed, within areas of high suitability, the most common land cover type (45%) is the ‘closed (>40%) needle-leaved evergreen forest (>5m)’ (i.e. coniferous forest), followed by the land cover types including croplands (36%). Contrary to the results obtained for Taurus ground squirrels, none of the bioclimatic variables clearly dominated over another one, suggesting that both temperature and precipitation (or the factors associated with these) are among the main drivers of the distribution patterns of coniferous forests. Indeed, at regional-to-global scales, climate exerts a dominant influence on the distribution patterns of plant functional types. However, smaller-scale variations in the distribution patterns may be controlled by smaller-scale features of the environment (e.g. soil and topography; Sykes & Haxeltine, 2013).

There is very limited overlap between Taurus ground squirrels and coniferous forests in areas of high suitability (Figure 3), suggesting that the climate envelopes of these species/communities are highly distinct. The most important variables in separating between these envelopes are temperature variables (stepwise discriminant analysis, results not shown). Taurus ground squirrels had an estimated niche breadth 1.7 times smaller than coniferous forests (0.230 vs. 0.397, respectively), suggesting that, as expected, high-altitude environments have a smaller subset of environmental variation than low-altitude ones (see also Table 1).

The high mountains rising steeply from the sea, the orientation of these mountains, especially in relation to the wind, and the prevailing southwest winds, carrying most of the rain, are the main factors influencing the climate, and therefore the vegetation, of the Mediterranean Taurus Mountains (World Wildlife Fund, 2019). In this region, precipitation mainly decreases from the south/southwest to the north/northeast (e.g. from the sea-facing side of Geyik Mountains to the inland areas), with increasing distance from the Mediterranean Sea, because these mountain ranges create a natural orographic barrier between the Mediterranean coast and the Anatolian plateau. However, temperature is mainly a function of altitude (Figure 6 and 7). The vegetation differs considerably between the lower and higher altitudes and between the sea- and inland-facing sides of the Mediterranean Taurus Mountains (e.g. Atalay & Mortan, 2006) mainly because of these gradients (this study). The results simply illustrate how these gradients have had a substantial role in shaping the distribution patterns of species/communities and therefore the biodiversity of these mountain ranges. Thus, particular sources of concern are the synergistic effects of future global climate change and anthropogenic impacts (e.g. transhumance, shifts in land-use systems, overgrazing) on high- and low-altitude species/communities (e.g. Taurus ground squirrels and coniferous forests, respectively) (for further discussion, see Gür et al., 2018).

**Figure 6.**
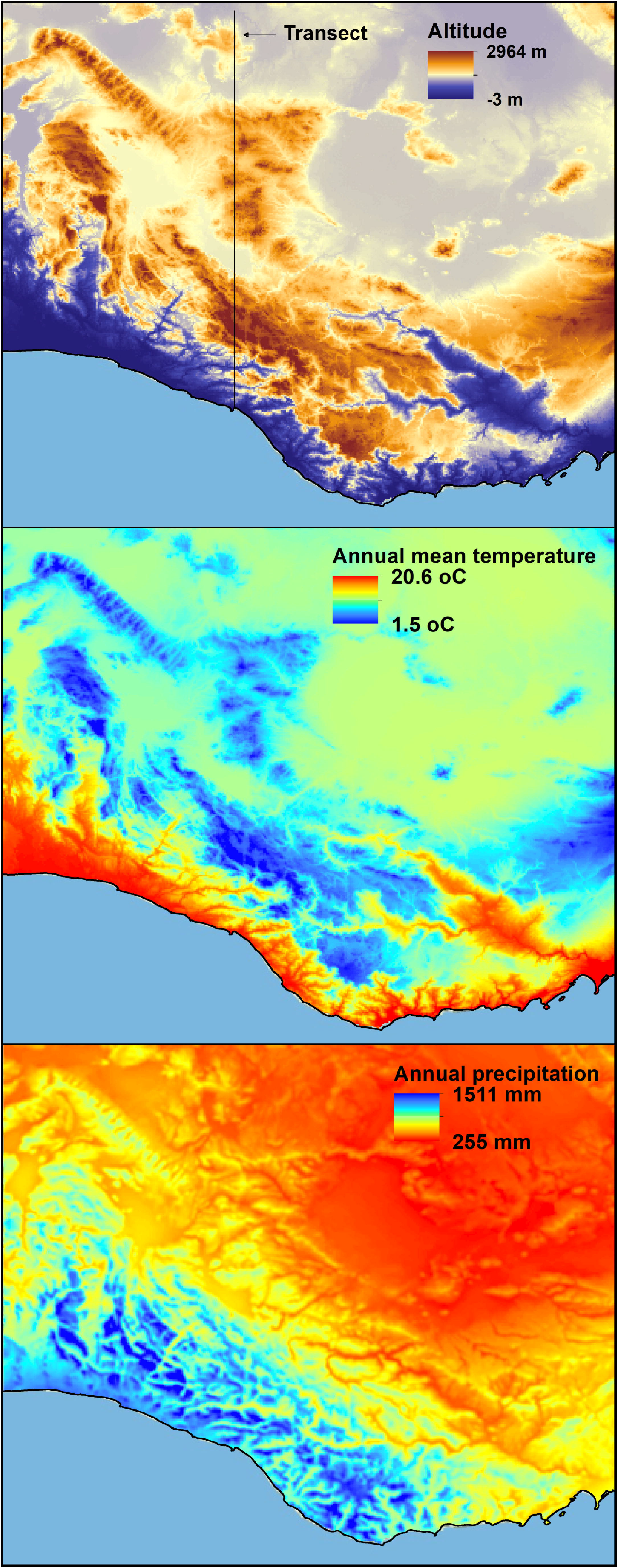
The spatial distributions of altitude, annual mean temperature, and annual precipitation in the study area. For the transect, see Figure 7.

**Figure 7.**
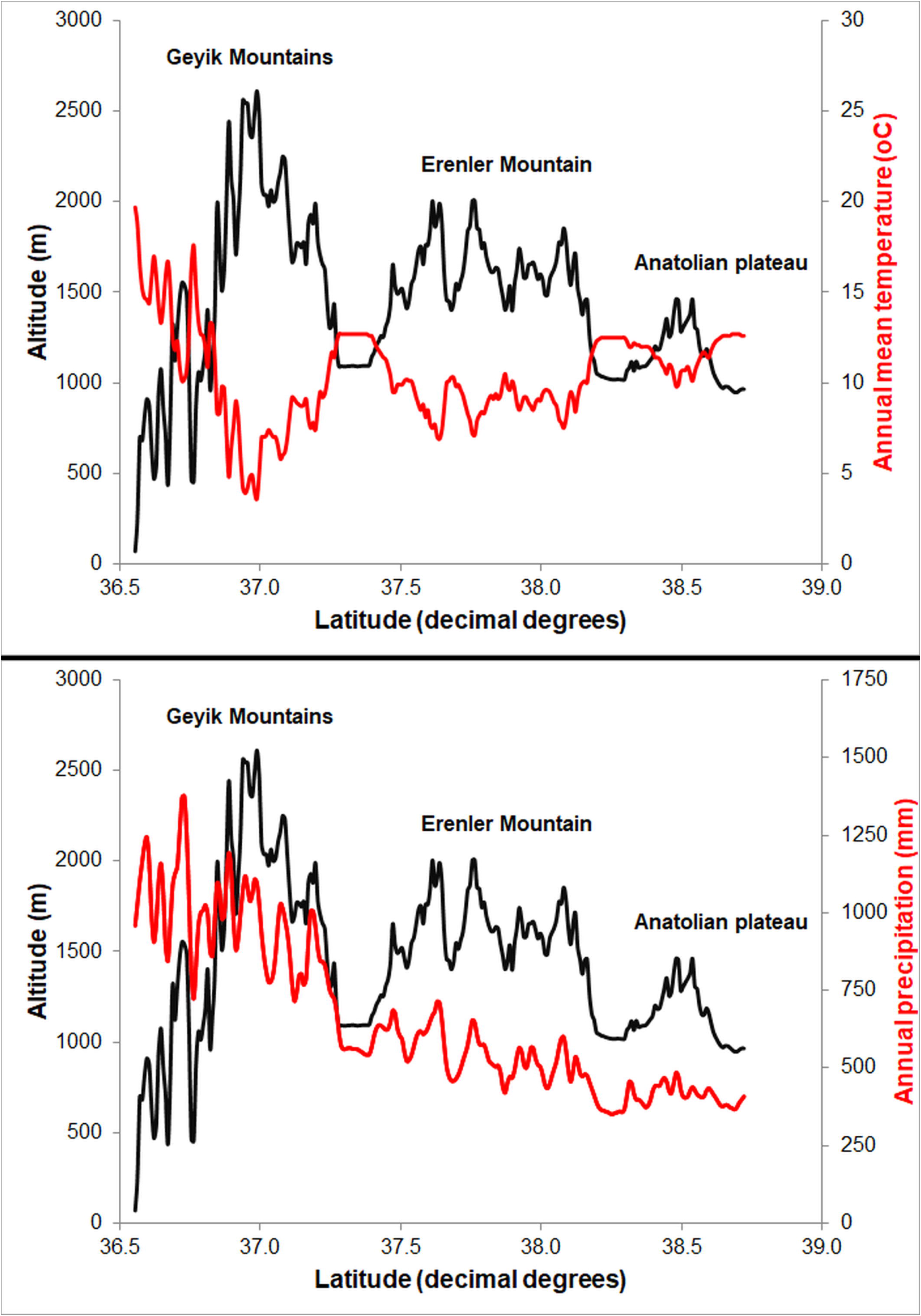
The spatial variations of altitude, annual mean temperature, and annual precipitation along a north-south transect (i.e. a black line shown at the top of Figure 6) in the study area.

## Acknowledgments

I would like to thank Çağatay Tavşanoğlu for providing helpful comments on the manuscript.

